# Green self-immolative polymer: molecular antenna to collect and propagate the signal for zymogen activation

**DOI:** 10.1101/2021.07.25.453687

**Authors:** Mireia Casanovas Montasell, Pere Monge, Sheiliza Carmali, Livia Mesquita Dias Loiola, Dante Guldbrandsen Andersen, Kaja Borup Løvschall, Ane Bretschneider Søgaard, Maria Merrild Kristensen, Jean Maurice Pütz, Alexander N. Zelikin

## Abstract

Chemical zymogens of three different types were established herein around protein cysteinome, in each case converting the protein thiol into a disulfide linkage: zero length Z_0_, polyethylene glycol based Z_PEG_, and Z_LA_ that features a fast-depolymerizing fuse polymer. The latter was a polydisulfide based on a naturally occurring water-soluble lipoic acid. Three zymogen designs were applied to cysteinyl proteases and a kinase and in each case, enzymatic activity was successfully masked in full and reactivated by small molecule reducing agents. However, only Z_LA_ could be reactivated by protein activators, demonstrating that the macromolecular fuse escapes the steric bulk created by the protein globule, collects activation signal in solution, and relays it to the enzyme active site. This afforded first-in-class chemical zymogens that are activated via protein-protein interactions. For Z_LA_, we also document a “chain transfer” bioconjugation mechanism and a unique zymogen exchange reaction between two proteins.

---

Self-immolative polymers (SIPs) are a cunning tool of polymer chemistry and materials science. ^1-8^ These polymers undergo triggered end-to-end decomposition when a stimulus is applied at a polymer chain end. SIPs are important for polymer recycling, ^9 10^ drug delivery, ^1, 11^ biosensing, ^12-13^ and materials science, ^14-18^ and hold immense potential for diverse areas of science and (bio)technology. Of the developed SIP, polydisulfides based on the derivatives of lipoic acid (LA PDS, Figure 1A) are unique in that these polymers are water soluble and to our knowledge are the only SIPs based on a naturally occurring compound.^19-22^ Currently, the prime utility of LA PDS is in the field of intracellular drug delivery, due to the capacity of a disulfide bond to undergo degradation upon cell entry. ^23-25 26^ Herein, we propose that the “green” characteristics of LA PDS make it highly attractive for a unique application, namely engineering a macromolecular fuse to collect and propagate chemical information between two proteins, to achieve activation of chemical zymogens (Figure 1B).

**Figure 1.**
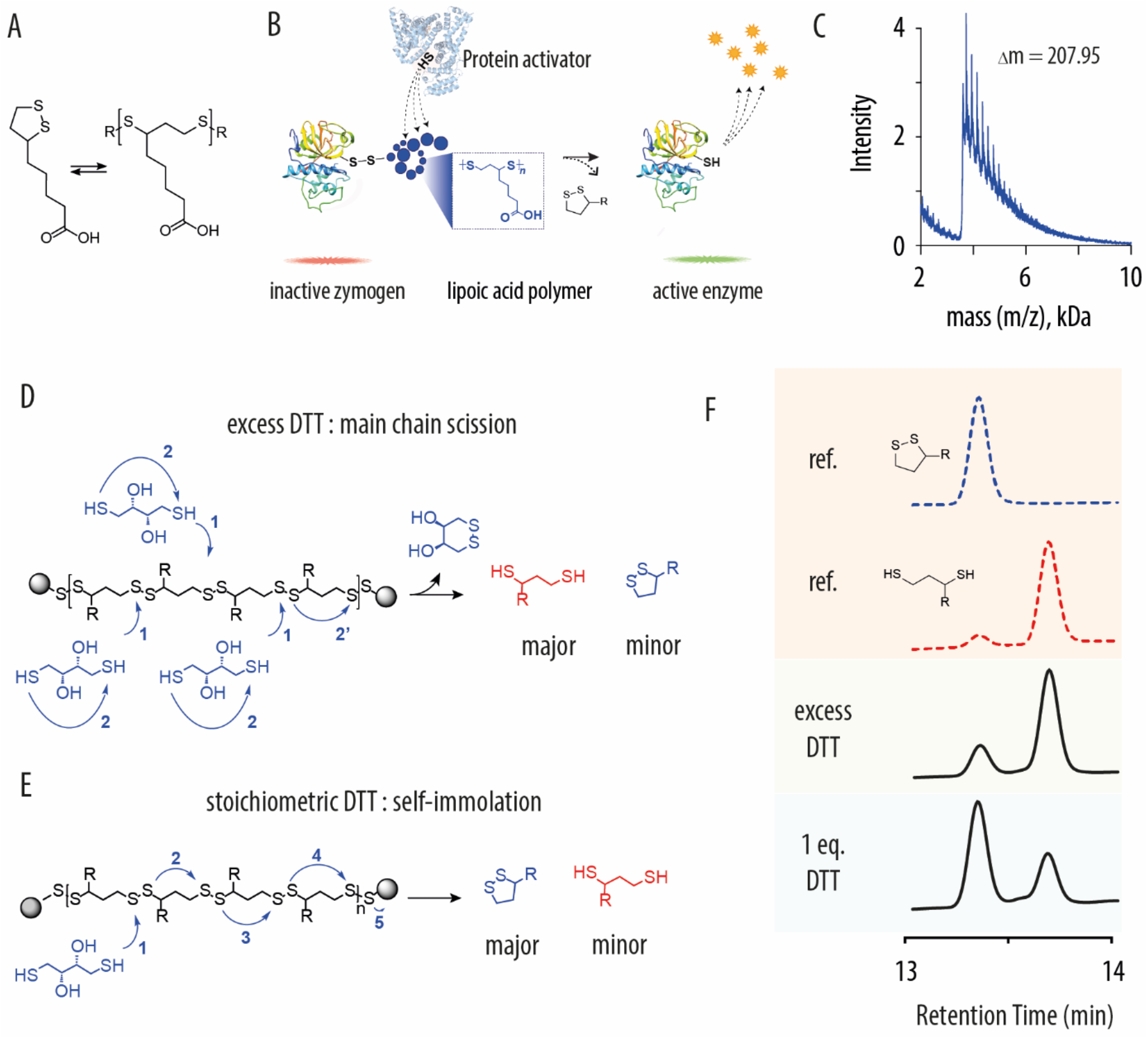
**(A)** Chemical formula of lipoic acid and schematic illustration of its reversible polymerization into a linear disulfide; **(B)** Schematic illustration of the proposed self-immolative antenna as a tool to collect activating chemical signal in solution and propagate it to the sterically hindered active site of an enzyme; **(C)** MALDI spectrum of the lipoic acid polymer that shows inter-peak spacing of 207.95 Da, in agreement with the structure of lipoic acid as the monomer unit; **(D**,**E)** schematic illustration of mechanisms of depolymerization for LA PDS via main chain scission (D) or self-immolation (E), depending on stoichiometry of added reducing agent; **(F)** reference HPLC traces of oxidized and reduced lipoic acid monomers, and experimental data on depolymerization of LA PDS in excess DTT and with equimolar DTT, illustrating main-chain scission or self-immolation as predominant mechanisms of depolymerization, depending on stoichiometry of reaction.

Mechanisms of reversible enzyme (de)activation in nature form the basis of signal transduction cascades and also lay behind the ability of cells and multicellular organisms to maintain the overall enzymatic homeostasis without wasteful, repetitive protein degradation and synthesis.^27-29^ Nature has developed robust mechanisms to perform zymogen-enzyme interconversion, most notably via the reversible (de)phosphorylation or via the irreversible proteolytic activation. In contrast, design of chemical, synthetic zymogens has proven to be highly challenging. First of all, it requires the identification of an essential amino acid in the protein sequence that can be modified so as to exert reversible masking of catalytic activity, and the identification of associated chemistry to achieve this. ^2, 30-32^ In this regard, the most promising opportunity is associated with the protein cysteinome: modification of cysteine thiols into mixed disulfides within the active site of cysteinyl protease^33^ or peptidase^2^, or catalytically nonessential thiol proximal to the active site in a kinase ^34^ has been shown to afford reversible enzyme deactivation. Nevertheless, compared to thousands of proteins that comprise the human cysteinome, and despite the growing importance of cysteinome inhibitors in medicinal chemistry, ^35^ synthetic zymogens built around the cysteine thiol are solitary. Furthermore, successful in their own right, in all cases, known synthetic zymogens have been activated by external small molecule chemical stimuli and not via proteinprotein interaction, as is the hallmark of biological enzyme-zymogen interconversions. This highlights the second, critical challenge in the zymogen design, namely the steric hindrance created by the protein globule, which protects the protein interior (cysteine thiols) from access by protein activators. ^33^

In this work, we address the fundamental challenge of developing chemical zymogens and specifically focus on the design of zymogens amenable for activation via protein-protein interactions. Towards the overall goal, we establish three types of chemical zymogens, in each case converting the protein thiol into a disulfide linkage: zero length zymogen Z_0_ (methyl disulfide-blocked essential thiol), polyethylene glycol based Z_PEG_, and zymogens featuring LA PDS as a fast depolymerizing fuse polymer, Z_LA_. The three designs are applied to two cysteinyl proteases and a kinase and investigated in terms of masking of enzymatic activity, recovery of catalysis by small molecule activators, and zymogen reactivation by protein activators. For the latter case, we also investigate the zymogen exchange reactions whereby the masking group is transferred from one protein to another. Results of this study establish broadly applicable zymogen design protocols and illustrate that only Z_LA_ afforded recovery of enzymatic activity by protein activators. This is because LA PDS aided to overcome the steric constraints exerted by the protein globule: the macromolecular fuse extends from a thiol in the protein globule into the solution bulk, wherein it interacts with the protein activators (Figure 1B). These results illustrate the successful transfer of chemical information from solution bulk to the enzyme active site by a macromolecular fuse and present, to our knowledge, the first example of a chemical zymogen reactivated using protein-activators.

The application of LA PDS as a macromolecular fuse to collect and propagate the signal requires an understanding of the polymer decomposition mechanism. We synthesized LA PDS via the thermally induced ring opening polymerization using iodoacetamide as a chain terminating reagent. Polymer composition was confirmed via MALDI (Figure 1C), which illustrated that the obtained polymers had lipoic acid as a monomer unit. Size exclusion chromatography (SEC) revealed typical molar masses of 20-40 kDa and dispersity indexes for the synthesized polymers between 1,2-1,6. Triggered polymer decomposition was monitored using HPLC which revealed that LA PDS degrades via two distinctly different mechanisms, depending on the stoichiometry of added reducing agent (Figure 1D-F). In excess dithiothreitol (DTT), polymer decomposition afforded reduced lipoic acid as the main product (Figure 1D,F), indicative of the main-chain scission of the polymer at multiple disulfide linkages. In contrast, one mole-equivalent DTT to polymer chains exerted a single point scission of the polymer backbone, and decomposition of LA PDS proceeded predominantly via the self-immolation mechanism, that is, through sequential ring-closing of monomer units (Figure 1E,F). Self-immolation is most desired for the envisioned use of LA PDS as a macromolecular fuse in that signal collection is propagated from the initiation point to the chain termini. Noteworthy, to our knowledge, there is only one, very recent prior example of a polymer that can degrade by two mechanisms, main chain scission or self-immolation, and kinetics of the latter process was measured in hours. ^36^ Uniquely, depolymerization of LA PDS was rapid and complete within minutes (Figure 1F), and initiation could occur at any disulfide bond in the backbone, not necessarily at the chain end, as is most typical for other SIP systems. ^4-5^

The zymogen design was first performed for a cysteinyl protease papain. The reaction between papain and LA PDS proceeded readily in their aqueous mixtures and afforded the protein-polymer conjugate (Figure 2A). This was confirmed using SEC, and in a typical experiment we observed that the 23 kDa protein was incorporated into the 39 kDa protein-polymer conjugate (Figure 2B). Isoelectric focusing gel electrophoresis indicated a change in the protein pI from over 8 (the protein does not enter the gel) to under 6, as would be expected for a conjugate with a polymeric acid (Figure 2C). From the standpoint of the mechanism, the reaction between the protein and the polymer appears to be rather unique in that the protein thiol performed a nucleophilic attack at the disulfide linkage within the polymer backbone. This is different from the “grafting to” coupling reactions^25 24^ and from the “grafting from” polymerization mechanisms ^2, 8, 23, 25^. The protein effectively “picked up” one part of the polymer chain, and bioconjugation occurred as a “chain transfer” reaction. This mechanism is routinely observed in nature (e.g. transferase activity on polypeptides) but to our knowledge has not been observed for synthetic macromolecules.

**Figure 2.**
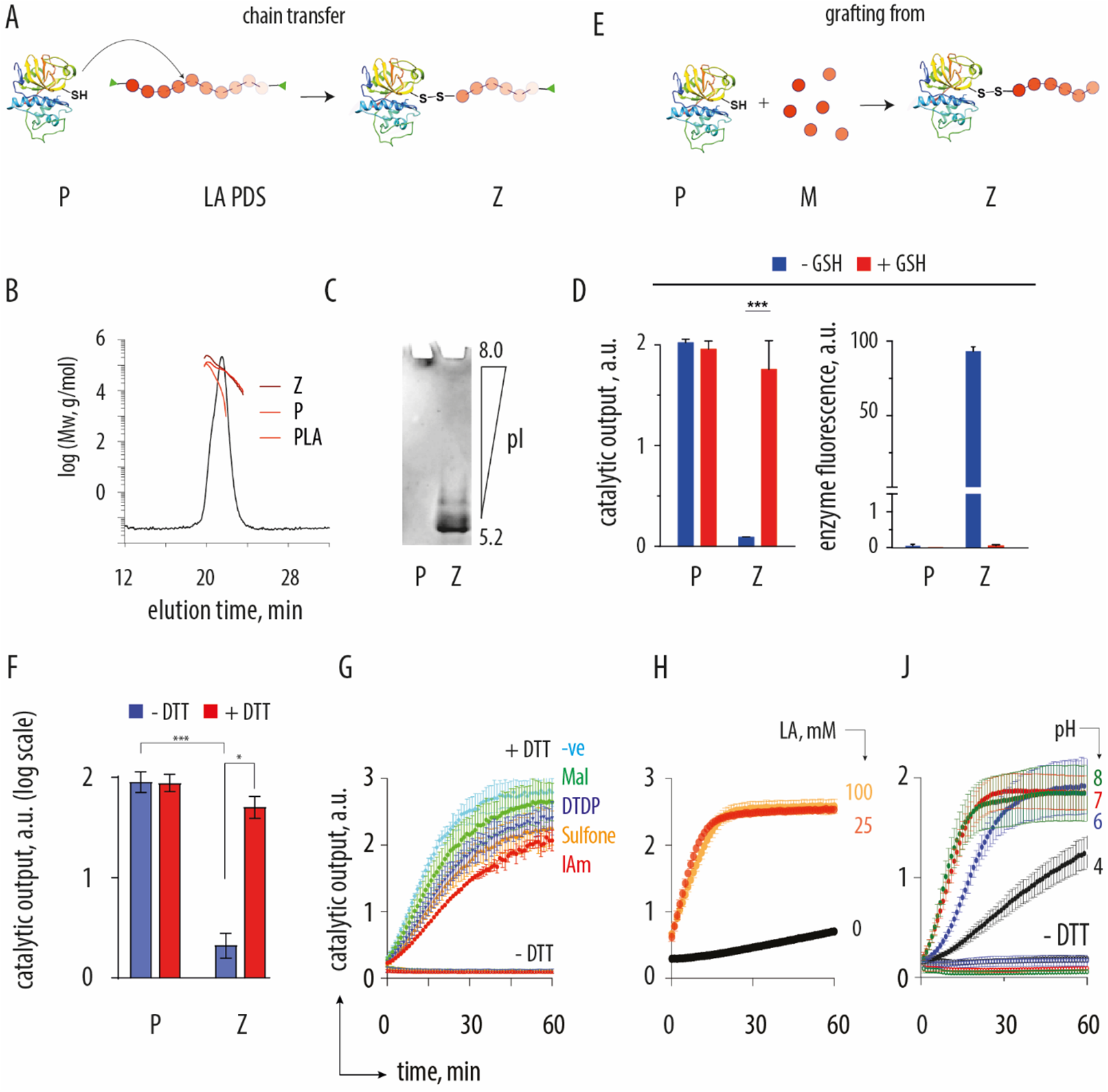
(**A**) Schematic illustration of the zymogen preparation reactions via the “chain transfer” mechanisms, P = protein, LA PDS = poly(lipoic acid), Z = zymogen; **(B)** SEC-MALS characterization of the papain – LA PDS conjugate; **(C)** isoelectric focusing gel electrophoresis analysis of papain and its conjugate with LA PDS; **(D)** Protein fluorescence and end-point measurement of enzymatic activity of papain and the LA PDS-containing zymogen whereby LA PDS has fluorescein as a terminal group; enzyme substrate : 50 µM N_α_-benzoyl-L-arginine-7-amido-4-methylcoumarin hydrochloride; [P, Z] = 1 µM; **(E)** Schematic illustration of the zymogen preparation reactions via the “grafting from” cryopolymerization route; P = protein, M = monomer; Z = zymogen; **(F)** end point measurements of enzymatic activity for P and Z derived thereof via cryopolymerization, illustrating successful deactivation of papain by LA PDS and reactivation of catalysis via polymer decomposition; (**G**) Kinetic data for enzymatic catalysis in solutions of zymogens obtained from papain via cryopolymerization of lipoic acid and post-treatment with thiol traps (IAm = iodoacetamide, DTDP = 2,2’ dithiodipyridine, Mal = 4-maleimidobutyric acid; Sulfone = phenyl vinyl sulfone; -ve = non-capped cryo-polymerization), upon optional reactivation with DTT; (**H**) enzymatic kinetics data in solutions of papain exposed to iodoacetamide with or without cryopolymerized LA; **(J)** pH dependence of kinetics of zymogen reactivation. Panel G: shown representative results from individual runs; panels D, F, H, J : shown are results from at least three independent experiments as mean ± S.D.

The thiol group in the active site of papain is indispensable for enzymatic activity of papain, and blocking this thiol should render the enzyme inactive.^33^ Indeed, in our hands, papain lost its enzymatic activity upon incubation with LA PDS, providing further proof of successful bioconjugation (Figure 2D). The polymer used in this experiment was synthesized using fluorescein iodoacetamide end-group capping, and its conjugate with papain correspondingly exhibited strong fluorescence. Addition of a reducing agent restored enzymatic activity of papain and at the same time resulted in the complete loss of protein fluorescence. This indicates degradation of LA PDS and release of the enzyme from its zymogen, and thus validates the zymogen design.

Zymogen synthesis was also performed using the recently introduced “grafting from” cryopolymerization^2^ of LA PDS using papain as an initiator (Figure 2E). Resulting products exhibited blocked catalytic activity, which was readily restored via the triggered depolymerization of the macromolecular fuse (Figure 2F). Cryopolymerization could be terminated using a range of thiol traps such as maleimide, dithiodipyridine, sulfone, or iodoacetamide, which offers further opportunities for advanced bioconjugation endeavors (Figure 2G). Surprisingly, non-quenched cryopolymerization also offered a zymogen preparation, most likely via oxidative coupling / chain growth termination. In each case zymogen was readily reactivated into the corresponding enzyme via polymer decomposition (Figure 2G). Noteworthy, LA PDS macromolecular fuse was essential in the zymogen design and the direct reaction between iodoacetamide (IAm) and papain afforded irreversible protein inhibition with a minimal catalytic output (Figure 2H). Finally, zymogen reactivation was found to be a pH-dependent process, being slowest at pH 4 and rather similar at pH 6 to 8, being consistent with the kinetics of the thiol-disulfide exchange (Figure 2J). The cryo-polymerization method of zymogen design proved to be advantageous in terms of zymogen purification and unless indicated otherwise was adopted for the syntheses of Z_LA_ in all experiments presented below.

To conduct a broader investigation of opportunities in the zymogen design and reactivation, two more classes of zymogens were synthesized for papain, namely the zero-length zymogen (Z_0_) and a PEG-based counterpart (Z_PEG_). In each case (as well as for the Z_LA_), the essential protein thiol was modified into a mixed disulfide functionality, methyl-for Z_0_ or a non-degradable 6 kDa polymer for Z_PEG._ Zymogens were analyzed for quenching and recovery of enzymatic activity via the fluorescence based read-out, and for composition via MALDI (Figure 3A). The latter analysis revealed the expected minimal change in molar mass for Z_0_ and a change of ∼5800 Da for Z_PEG_, consistent with the addition of a single PEG chain to papain, as well as the broad peak for Z_LA_ with masses higher than papain. We note that MALDI analyses are typically accompanied with the in-flight decomposition of the labile disulfide functionality (routinely seen for antibodies), which explains the observed pristine papain in Z_PEG_ and only moderate molar mass increase for Z_LA_. Nevertheless, for each zymogen, enzymatic activity of the protein was masked to negligible levels, indicating quantitative modification of the essential thiol into mixed disulfides. Addition of a small molecule reducing agent (DTT) readily restored enzymatic activity, validating that each of the three zymogen designs was successful. The three robust protocols for the zymogens design were readily applied to another cysteinyl protease, bromelain: for each zymogen (Z_0_, Z_PEG_, Z_LA_) we achieved exhaustive suppression of enzymatic activity, which was restored in the presence of reducing agents (Figure 3A).

**Figure 3.**
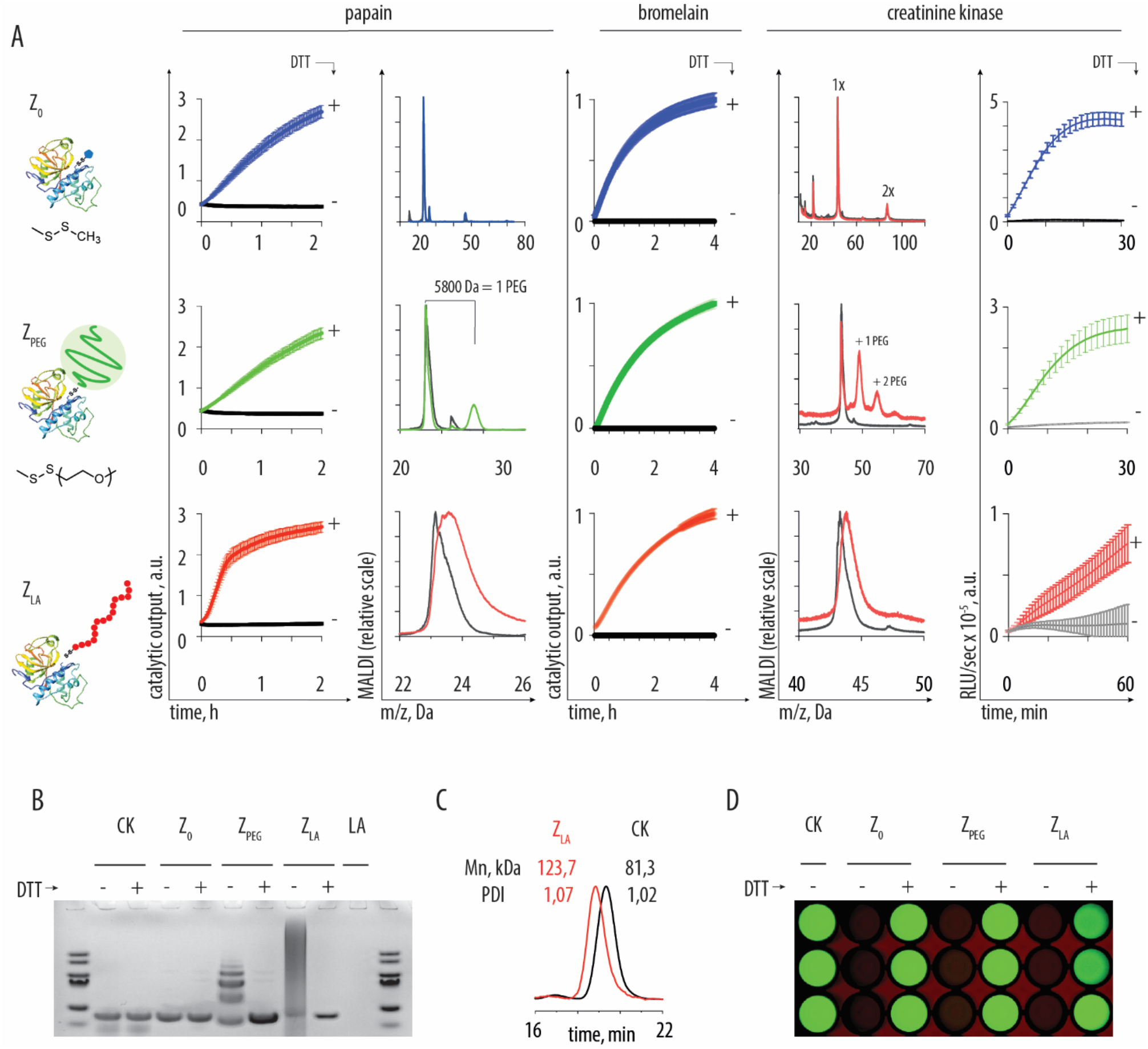
(A) Schematic illustration of the zymogens design, the chemical formulas of the corresponding disulfides, the reactivation data, and the MALDI characterization data for the zero-length zymogen Z_0_, PEG-based zymogen Z_PEG_, and the zymogen with the macromolecular fuse Z_LA_; for experimental condition see Supporting Information; shown results are based on three independent experiments (starting with zymogen syntheses); (B) gel electrophoresis characterization of Z_0_, Z_PEG_, and Z_LA_ synthesized using creatine kinase; (C) SEC-MALS characterization of the creatine kinase Z_LA_; (D) luminescence end-point imaging of the coupled kinase / luciferase assay illustrating activity of CK masked in the composition of the three zymogens and recovered with the use of DTT as a reducing agent.

To broaden the scope of the chemical zymogen methodology even further, the three types of zymogens (Z_0_, Z_PEG,_ Z_LA_) were also synthesized for a member of the kinase protein family, namely creatine kinase (CK). This protein has 4 cysteine thiols, one of which is found in the proximity of the active site but is nevertheless non-essential for catalysis. Gel electrophoresis and MALDI analyses confirmed addition of multiple PEG chains per protein for Z_PEG_ (Figure 3A,B), and the increase in molar mass for Z_LA_ over pristine kinase. SEC analysis for Z_LA_ confirmed protein-polymer conjugation, pristine kinase eluting as a dimer species with observed molar mass close to the theoretical value of 85 kDa and the conjugate revealing a molar mass of over 120 kDa (Figure 3C). For each zymogen composition, gel electrophoresis confirmed that the protein was released from its conjugates upon addition of DTT (Figure 3B). By mechanism of action, CK uses creatine phosphate as a substrate and catalyzes conversion of ADP into ATP. Taking advantage of this, activity of the kinase was quantified through a secondary read-out, via quantification of ATP in a luciferase-based assay. In each case, solutions of zymogens and luciferase (supplemented with ADP, creatine phosphate, and luciferin) exhibited minor luminescence (Figure 3A). Upon addition of DTT, solutions revealed progressively higher luminescence with time, indicative of the build-up of concentration of ATP and illustrating recovery of the kinase from its zymogens. Imaging of the reaction wells provided visual confirmation of the zymogen design and reactivation for Z_0_, Z_PEG,_ Z_LA_ (Figure 3D).

Taken together, results in Figure 3 illustrate the successful design of three novel chemical zymogens, applied to three different proteins. We believe that the design methodology presented herein is applicable to a diverse range of proteins that comprise cysteinome, including the proteins with thiols that are essential for catalytic activity and the thiols that are non-essential for catalysis, and providing structural diversity (zero length, non-degradable protein, fast-depolymerizing macromolecular fuse) to suit particular applications in biotechnology or biomedicine.

For protein activators, we observed a drastic difference in the zymogen reactivation phenomena between the three zymogens. To demonstrate this, we used papain-based zymogens. As protein activators, we used lysozyme, creatine kinase, pyruvate kinases (type VII and II, from rabbit muscle), and transglutaminase. For Z_0_ and Z_PEG_, we observed negligible catalysis on a fluorogenic papain substrate in the presence of protein activators, indicating negligible zymogen reactivation (Figure 4A). This was also true for Z_LA_ mixed with lysozyme, a thiol-free protein taken as a negative control. In stark contrast, for the thiol containing proteins used as protein activators for Z_LA_, we observed significant increase in solution fluorescence, which reports on the zymogen reactivation. For creatine kinase and pyruvate kinase VII, this was observed at sub-micromolar Z_LA_ content, whereas for pyruvate kinase II and transglutaminase it required a higher Z_LA_ concentration of at least 3 µM. In all cases, enzymatic activity was statistically significant compared to both Z_0_ and Z_PEG_ in the presence of the same protein activator. Connectivity at the papain Cys thiol for each zymogen (Z_0_, Z_PEG,_ Z_LA_) is a disulfide linkage and zymogen designs only differ in steric aspects. Zymogen reactivation for Z_LA_ was therefore achieved specifically owing to the macromolecular fuse, which extended from the protein essential thiol into solution bulk to interact with protein activators and thereafter propagate the activation signal to the enzyme active site via depolymerization. Zymogen reactivation occurred within minutes, owing to the fast depolymerization kinetics of LA PDS.

**Figure 4.**
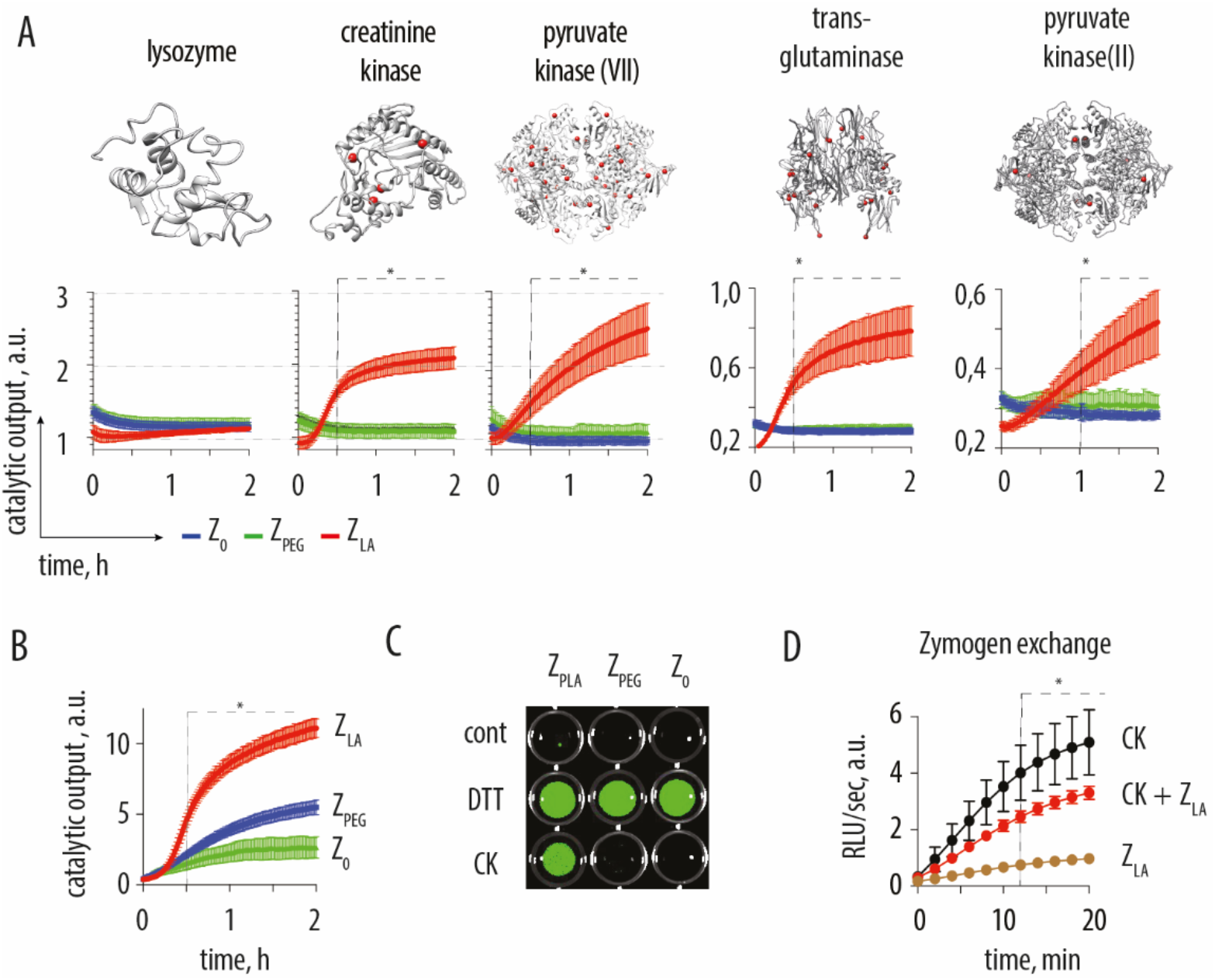
**(A)** Kinetics of zymogen reactivation using protein activators in 25mM borate buffer pH8 (+25mM NaCl, 1mM EDTA); for reactivation with creatine kinase and pyruvate kinase VII, concentrations were Z_0_, Z_PEG_ = 1 µM, Z_LA_ = 0,3 µM, papain substrate concentration 10 µM; for reactivation with transglutaminase and pyruvate kinase II: Z_0_, Z_PE_G = 10 µM, Z_LA_ = 3 µM, papain substrate concentration 10 µM; **(B)** kinetic measurements for reactivation of papain zymogens using creatine kinase, recorded using albumin (blocked Cys-34, labelled with fluorescein isocyanate over the level of self-quenching) as a protease substrate; **(C)** end point luminescence imaging illustrating that papain activity is masked by the three zymogens and revealed by DTT for Z_0_, Z_PEG_ and Z_LA_, whereas recovery with the protein activator is possible only for Z_LA_; **(D)** zymogen exchange reaction illustrated by a decrease of the kinase activity (recorded via the coupled kinase / luciferase assay) upon addition of papain Z_LA_. In Panels A, B, D: data shown are an average of three independent experiments (starting with independent zymogen syntheses) shown as mean ± st.dev., statistical analysis was carried via the two-way ANOVA test with statistical significance indicating the time-point at which Z_LA_ reactivation becomes statistically significant compared to both, Z_0_ and Z_PEG_ (in panels A,B) or compared to the activity of creatine kinase.

To increase the biological relevance of the data, we realized a three-enzyme cascade, using albumin as a substrate to quantify proteolytic activity of papain. Specifically, we used creatine kinase as an activator and attempted to release papain from Z_0,_ Z_PEG_ and/or Z_LA_ and in doing so to activate papain for activity at a natural substrate, albumin (blocked at Cys-34 and fluorescently labelled over the self-quenching level). This three-protein cascade was quantitatively analyzed using fluorescence read-out (Figure 4B) and visualized on an imaging plate reader (Figure 4C). These data exemplify the overall results of this study and show that Z_0_, Z_PEG,_ Z_LA_ compositions are equally powerful as zymogen design chemistries in masking protein activity, but only Z_LA_ equipped with a macromolecular fuse can be reactivated through an interaction with another protein, an activating kinase.

Kinase activity as a protein activator for papain Z_LA_ suggests the occurrence of a thiol-disulfide exchange reaction between the kinase thiol(s) and the papain-polymer conjugate. This reaction liberates papain from its zymogen (Figure 3,4) and at the same time should therefore produce the kinase conjugate with LA PDS. In other words, by activating the papain Z_LA_, kinase protein should be converted from an active enzyme to the corresponding zymogen. To validate this, we quantified the kinase activity via the coupled luciferase read-out. Addition of papain Z_LA_ led to a statistically significant decrease in the kinase activity (Figure 4D), indicating conjugation of CK into the corresponding Z_LA_. This evidence validates the occurrence of the “chain transfer” event wherein part of the LA PDS is transferred from one protein to another, in what is to our knowledge the first observation of a zymogen exchange reaction. LA PDS is therefore a unique tool for macromolecular science and chemical biology, to synthesize chemical zymogens based on the protein cysteinome, to achieve triggered recovery of the enzyme activity, and to achieve zymogen exchange reactions.

## Conclusions

Taken together, results of this study present i) the development of chemical zymogens around the protein cysteinome, ii) the development of chemical means to achieve zymogen reactivation using protein activators, and iii) the engineering of means to establish zymogen exchange reactions. Three classes of zymogens were developed in this work: the zero-length Z_0_, Z_PEG_ based a non-degradable polymer, and Z_LA_ that features a fast depolymerizing self-immolative polymer. These were applied to proteins of different classes (proteases, kinase). In each case, enzymatic activity was successfully masked in full and reactivated by small molecule reducing agents, indicating successful zymogen design. Protein activators could only restore activity in the case of Z_LA_, highlighting the critical need for the macromolecular fuse to extend into the solution bulk, collect the activation signal, and propagate it to the protein interior via depolymerization. Significant novelty of this work also includes the “chain transfer” bioconjugation reaction between a protein and a polymer, and the observed “zymogen exchange” reaction between two enzymes.

## Supporting information

Experimental section

## Acknowledgements

Authors acknowledge financial support from the Independent Research Fund Denmark (DFF FNU Grant No 0135-00162B, to ANZ), the Novo Nordisk Foundation (grant No NNF20OC0062131, to ANZ), the Carlsberg Foundation (Grant No CF19-0275, to ANZ), and the Lundbeck Foundation (grant No R287-2018-1117, to SC).

