## Supplementary material for "Green self-immolative polymer: molecular antenna to collect and propagate the signal for zymogen activation": Experimental section

1. Department of Chemistry and iNano Interdisciplinary Nanoscience Centre
   Aarhus University
   Aarhus C 8000
   Denmark

2. School of Pharmacy, Queen's University Belfast, United Kingdom

### 1. General Materials and Methods

All chemicals and proteins, unless stated otherwise, were purchased from Sigma Aldrich and used without purification. Deuterated solvents were supplied from Euriso-Top. Ultrapure water was dispensed from MilliQ Direct 8 (Millipore) [18.2 MΩ • cm].

**Nuclear Magnetic Resonance** (NMR) spectra were recorded on a Varian Mercury 400 MHz spectrometer, running at 400 MHz. Chemical shifts (δ) are reported in ppm relative to the residual solvent.

**Matrix Assisted Laser Desorption Ionization Time-of-Flight Mass Spectrometry (MALDI-TOF-MS)** were recorded using a Bruker Autoflex II MS with nitrogen laser (337 nm) in linear positive 20-200 kDa mode. At least 100 laser shots covering the complete spot were accumulated for each spectrum. For molecular weight determination, sinapinic acid (20 mg/mL) in 50 % acetonitrile with 0.1 % trifluoroacetic acid was used as matrix. Sample solution (0.1 to 1.0 mg/mL) was mixed with an equal volume of matrix and 4 μL of the resulting mixture was loaded onto a ground steel target plate and allowed to dry for cocrystallization.

**Size-Exclusion Chromatography with Multi-Angle Light Scattering (SEC-MALS)** was performed on a system containing an Agilent 1260 Infinity II Isocratic HPLC Pump, a Wyatt miniDAWN 3-angle static light scattering detector, an Agilent 1260 Infinity II VWD UV-vis detector, and a Shodex RI 501 refractive index detector. The system was equipped with a Superdex 200 Increase 10/300 GL column from GE Healthcare with a length of 300 mm, an internal diameter of 10 mm, and 8.6 µm particle size. This provided an effective molecular weight range of 10.000-600.000. The eluent was 0.01 M PBS with 10% methanol and it ran at a temperature of 30°C and a flow rate of 0.75 mL/min. Molar mass analyses were conducted using ASTRA® Software Basic.

**Reverse-phase HPLC** analyses were performed on an Agilent apparatus equipped with a C18 column with 2.7 µm particles, a length of 150 mm and an internal diameter of 3.0 mm from Supelco Analytical. HPLC mobile phase A was ultrapure $H_2O$ supplemented with 0.1 % TFA (v/v) and mobile phase B acetonitrile supplemented with 0.1 % TFA (v/v). Elution was performed starting with solvent B 5% to B 100% over 15 min, hold B 100% for 4 min at T = 40°C and a flow rate of 0.4 mL/min. Detection was performed by UV detector (220 nm and 280 nm).

**Fluorescence and luminescence measurements** were recorded on an Enspire 2300 Multilabel Reader (Perkin Elmer®) or a SynergyH1 microplate reader (BioTek®)

#### 2. Preparation of zymogens

##### 2.1 Ring-Opening Polymerization (ROP) of Lipoic Acid end-capped with 6-iodoacetamide-fluorescein

In a typical experiment, under argon atmosphere, lipoic acid (300.6 mg, 1.46 mmol) and 6-iodoacetamid-fluorescein (7.5 mg, 0.0146 mmol) were added to a flame-dried 25 mL pear-shaped glass flask. The mixture was heated to 80 °C for 2 hours. The resulting material was dissolved in 10 mL of 0.5 M sodium hydrogen carbonate buffer (pH 8.5), purified by dialysis (MWCO 3.5 kDa), and lyophilized to afford poly (lipoic acid) – fluorescein capped as an orange product (yield 162.9 mg, 54 %).

$^{1}$H NMR (400 MHz, Deuterium Oxide) δ 8.16 – 7.57 (m, 6H), 7.43 – 7.03 (m, 4H), 6.83 – 6.25 (m, 8H), 3.61 – 3.49 (m, 4H), 2.77 (s, 325H), 2.09 (s, 229H), 1.93 (s, 220H), 1.53 (s, 218H), 1.44 (s, 235H), 1.32 (s, 225H).

SEC: DP (NMR) 108, $M_N$ (NMR): 23 kDa.

##### 2.2 Reduction of papain and bromelain

Papain and bromelain, as cysteine proteases, are commercially provided as partially inactivated enzymes. Activation is accomplished by incubation with a thiol-containing compound or a reducing agent. In a typical experiment, to a solution of papain (5 mg/mL, 0.17 mM, 0.5 mL) in 50 mM phosphate buffer pH 6.8 with 10 mM EDTA was added tris (2-carboxyethyl) phosphine hydrochloride (TCEP, 1.1 mM, 5 equiv). The reaction was incubated at 37°C for 1 h. After this time, TCEP was removed by gel filtration (NAP-5, GE Healthcare) in the same buffer and concentrated using spin filtration (MWCO 3 kDa).

##### 2.3 Preparation of PEG-S-S-TP

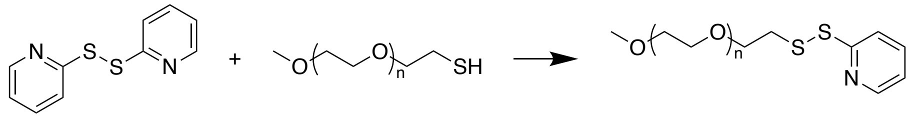

A solution was prepared of 2,2'-dithiodipyridine (45.8 mg, 4.0 equiv.) and acetic acid (1.5 µL, 0.5 equiv.) in methanol (7.5 mL) under an atmosphere of argon. MeO-PEG-SH-6000 (312.0 mg, 1.0 equiv.) was dissolved in a mixture of methanol:acetic buffer 50 mM (pH = 4.5) 1:1 v:v (2 mL), purged with argon and added dropwise to the reaction mixture. The reaction was left stirring at room temperature for 21 hours. The product was precipitated twice into cold diethyl ether followed by centrifugation. The precipitated product was obtained by filtration followed by solvent removal *in vacuo* to yield PEG-S-S-TP (176.0 mg, 53%) as a white solid.

**$^{1}$H-NMR** (400 MHz, Methylene Chloride-$d_2$) δ 8.45-8.41 (m, 1H), 7.83-7.78 (m, 1H), 7.71-7.65 (m, 1H), 7.13-7.07 (m, 1H), 3.77 (t, $J$ = 5.2 Hz, 2H), 3.71-3.41 (m, 144H), 3.34 (s, 3H), 3.00 (t, $J$ = 6.0 Hz, 2H).

##### 2.4 General protocol for zero length zymogens

Reduced papain, bromelain ( both at 2 mg/mL) or creatine kinase (1 mg/mL) were reacted with S-methyl methanethiosulfonate (MMTS, 500 equiv.) in sodium phosphate buffer (50 mM, 10 mM EDTA, pH 6.8). The solution was stirred at 4º C overnight and the proteins were purified / buffer exchanged to sodium acetate buffer (2.5 mM, pH 4.5, 50 mM NaCl) using Amicon centrifugal filters (MWCO 3 kDa)

##### 2.5 General protocol for PEG-based zymogens

Reduced papain and bromelain (100 μM) or creatine kinase (25 μM) were combined with 10 eq. of PEG-TP in acetate buffer (2.5 mM, pH 4.5, 50 mM NaCl) and incubated at 37º C for 1 hour. After this time, the crude sample was purified by gel filtration (NAP-5) in the same buffer.

##### 2.6 General "chain transfer" protocol for preparation of LA PDS zymogens

Reduced papain or bromelain (2 mg/mL) and excess LA PDS (10×, by weight) were dissolved in sodium phosphate buffer (50 mM, 10 mM EDTA, pH 6.8). The solution was stirred at 37 °C for 2 h. After this time, the crude reactions were purified by Amicon centrifugal filtration in the same buffer.

##### 2.7 General "grafting from" cryopolymerization protocol for preparation of LA PDS zymogens

Reduced papain (0.5 mg/mL) or creatine kinase (1 mg/mL) were dissolved in solutions containing 100 or 25 mM of LA in 0.2 M $NaHCO_3$ respectively, and 1 mL of these reaction solutions was kept in the freezer at -20º C for 2 hours. After this time, 200 μL of a 0.1 M solution of iodoacetamide in MQ water was added to the frozen reactions and left stirring at room temperature for 30 minutes. The zymogens were purified by gel filtration (CentriPure P10) in MQ water.

#### 3. Experimental protocols

##### 3.1 RP-HPLC analysis of poly (lipoic acid) degradation using Dithiothreitol (DTT)

Poly (lipoic acid) was incubated at room temperature in the presence and absence of dithiothreitol in 50 mM phosphate buffer, pH 8.0. At specific time points (t = 5 min and 24 h), reactions were analysed by RP-HPLC. Lipoic acid (0.10 mg/mL, 0.5 μmol) incubated under the same conditions served as controls.

##### 3.2 Isoelectric focusing

An isoelectric focusing (IEF) gel was performed to analyse the formation of papain-LA PDS conjugate. Samples were prepared of papain-LA PDS conjugate (0.8 g/L) and native papain (20 g/L) in 20 mM phosphate buffer, pH 6.8 with 1 mM EDTA. The samples were mixed with sample buffer (Novex® IEF Sample Buffer pH 3-10) in a 1:1 ratio and 10 μL of the solution was loaded to the gel (Novex™ pH 3-10 IEF protein gel). The gel ran for 1 hour at 100 V, 1 hour at 200 V and 30 minutes at 300 V at 4°C in Novex™ IEF Cathode Buffer pH 3-10 in the upper chamber and Novex™ IEF Anode Buffer in the lower chamber. The gel was washed in 1% AcOH in pure water for 2 hours (change of solution every 15 minutes) followed by staining with comassie blue (Invitrogen, LC6065).

##### 3.3 Reactivation of papain LA PDS zymogens

Solutions of 1 μM of papain and papain LA PDS zymogens prepared by either chain transfer or cryopolymzerization methods were incubated with 50 μM Nα-benzoyl-L-arginine-7-amido-4-methylcoumarin phosphate buffer (50 mM, 10 mM EDTA, pH 6.8). To each sample was added GSH to 10 mM or just buffer as a control with final reaction volumes of 100 μL. After 60 minutes at 37° C substrate hydrolysis was measured by recording fluorescence at λex/λem 370/460 nm.

##### 3.4 Reversible fluorescence labelling of papain LA PDS fluorescein zymogen

Papain-LA PDS fluorescein (1 mg/mL) was incubated with 10 mM GSH in 50 mM phosphate buffer, pH 6.8 with 10 mM EDTA for 60 min at 37 °C. After this time, the reaction mixture was purified by Amicon centrifugal filtration (MWCO 3 kDa). Native papain, papain-LA PDS fluorescein and purified papain-LA PDS fluorescein after incubation with GSH were separately measured for fluorescence from LA PDS conjugation ($\lambda_{ex}/\lambda_{em}$ 490/520 nm) with a temperature-controlled plate holder at 37 °C. Measured protein samples were assayed at 1 μM concentration in 50 mM phosphate buffer, pH 6.8 with 10 mM EDTA.

##### 3.5 Effect of different polymerization quenchers on LA PDS zymogen reactivation

LA PDS zymogens were prepared by cryopolymerization as described in section 2.7, but changing the quenching reagent used after polymerization. The quenchers used were: iodoacetamide, 2,2'dithiodipyridine, 4-maleimidobutyric acid and phenyl vinyl sulfone in concentrations of 0.1 M. An extra reaction in the absence of quencher was also prepared. After purification, equal volumes of each reaction were diluted in borate buffer (25 mM, pH 8.0) to a final protein concentration of 2 μM, combined with 5 μM of Nα-benzoyl-L-arginine-7-amido-4-methylcoumarin. Catalytic activity in the presence or absence of DTT (2 mM) was analysed by substrate hydrolysis in 100 μL reaction volumes as detected by fluorescence increase at $\lambda_{ex}/\lambda_{em}$ 370/460 nm at 1 minute intervals for 2 hours at 37° C in a plate reader.

##### 3.6 Effect of LA concentration during cryopolymerization

LA PDS zymogens of papain were prepared by cryopolymerization as described in section 2.7, but with changing concentrations of LA during the reaction to 100 or 25 mM and also a control without LA. After quenching of the reactions with iodoacetamide and purification, equal volumes of each reaction were diluted in borate buffer (25 mM, pH 8.0) to 0.3 μM protein (as determined by UV absorbance at 280 nm), combined with 5 μM of Nα-benzoyl-L-arginine-7-amido-4-methylcoumarin. Catalytic activity in the presence and absence of DTT (2 mM) was analysed by substrate hydrolysis in 100 μL reaction volumes as detected by fluorescence increase at $\lambda_{ex}/\lambda_{em}$ 370/460 nm at 1 minute intervals for 2 hours at $37^{o}$ C in a plate reader. These experiments were performed in 3 independent replicates (independent zymogen syntheses).

##### 3.7 Effect of pH on the reactivation of papain LA PDS zymogens

Papain LA PDS zymogen prepared by copolymerization method as in section 2.7 was dissolved at 5 μM and combined with 50 μM Nα-benzoyl-L-arginine-7-amido-4-methylcoumarin in different buffers with the addition of 10 mM DTT. Samples without DTT were prepared as controls. The different buffers were: formic acid ( 20 mM, pH 4.0), 2-(N-morpholino)ethanesulfonic acid (20 mM, pH 6.0), phosphate buffer (20 mM, pH 7.0) and borate buffer (25 mM, pH 8.0). Substrate hydrolysis was measured by recording fluorescence at $\lambda_{ex}/\lambda_{em}$ 370/460 nm at 1 minute intervals for 2 hours at $37^{o}$ C in a plate reader. These experiments were performed in 3 independent replicates (independent zymogen syntheses).

##### 3.8 Reactivation studies of papain and bromelain zymogens

Reactivation of different zymogens ($Z_0$, $Z_{PEG}$ and $Z_{LA}$) of papain and bromelain was studied by monitoring fluorescence increase upon hydrolysis of Nα-benzoyl-L-arginine-7-amido-4-methylcoumarin. Briefly, solutions containing papain zymogens ($Z_0$, $Z_{PEG}$ = 1 μM, $Z_{LA}$ = 0.3 μM as determined by UV absorbance at 280 nm) or bromelain zymogens ($Z_0$, $Z_{PEG}$, $Z_{LA}$ = 5 μM, as determined by UV absorbance at 280 nm) were combined with 10 μM of substrate and DTT (2 mM) in borate buffer (25 mM, pH 8.0) in a volume of 100 μL. Samples without DTT were used as controls. Fluorescence ($\lambda_{ex}/\lambda_{em}$ 370/460 nm) was monitored at 1 min intervals at $37^{o}$ C in a plate reader. These experiments were performed in 3 independent replicates (independent zymogen syntheses).

##### 3.9 Reactivation studies of creatine kinase zymogens

Reactivation of different zymogens ($Z_0$, $Z_{PEG}$ and $Z_{LA}$) of creatine kinase (CK) was quantified via a coupled bi-enzymatic assay. Briefly, unmodified CK or zymogens were diluted to 100 nM (as estimated by a protein assay with fluorescamine[2] using CK as reference) in Tris-HCl buffer (20 mM, pH 8.0) in a white micro plate. Then DTT was added, and quickly after, a solution containing ADP, creatine phosphate (CP), luciferin and Cell-Titer Glo® 2.0 Cell Viability Assay luciferase (Promega) was added to the wells. Luminescence was monitored at $37^{o}$ C in regular time intervals over 1 hour (RLU integrated every 2s) in a plate reader. Final volume in every well was 100 μL and concentration of reagents was as follows: DTT 20 μM, ADP, CP and luciferin 1 mM, Cell-Titer Glo® 2.0 1:50 dilution (v/v). Controls included samples of each zymogen/protein without DTT, CK without CP and samples without CK or zymogen. At the end point, luminescence was also imaged inside an ImageQuant™ LAS 4000 camera system (GE Healthcare) with 3 second exposure. These experiments were performed in 3 independent replicates (independent zymogen syntheses).

##### 3.10 Gel electrophoresis of creatine kinase zymogens

Samples of 2 μg of creatine kinase and creatine kinase zymogens were prepared in miliQ water supplemented with LDS buffer (Pierce, 1:4 final dilution) and NuPAGE™ sample reducing agent (in a final dilution of 1:8). For each protein or zymogen, a sample without reducing agent was also prepared. Samples were incubated 20 min at 37° C and subsequently 10 μL were loaded into a NuPAGE™ gel (4 to 12% bis-tris, 1.0 mm) and run for 50 minutes in MOPS buffer (Invitrogen NP0004) at 150 V and later stained with coomassie blue (Invitrogen, LC6065).

##### 3.11 Reactivation of papain zymogens with different protein activators

Reactivation of different zymogens ($Z_0$, $Z_{PEG}$ and $Z_{LA}$) of papain was studied by monitoring fluorescence increase upon hydrolysis of Nα-benzoyl-L-arginine-7-amido-4-methylcoumarin. Briefly, solutions containing papain zymogens ($Z_0$, $Z_{PEG}$ = 1 μM, $Z_{LA}$ = 0.3 μM as determined by UV absorbance at 280 nm) were combined with 10 μM of substrate and creatine kinase (CK) or pyruvate kinase VII at 1 μM in borate buffer (25 mM, pH 8.0). Reactivation was also performed with transglutaminase and pyruvate kinase II at 1 μM, but with a 10-fold increase in zymogen content ($Z_0$, $Z_{PEG}$ = 10 μM, $Z_{LA}$ = 3 μM). Fluorescence ($\lambda_{ex}/\lambda_{em}$ 370/460 nm) was monitored at 1 min intervals at 37° C in a plate reader. All well volumes were set to 100 μL and samples with DTT (1 mM) or without any activator were prepared as positive and negative controls respectively. These experiments were performed in 3 independent replicates (independent zymogen syntheses).

##### 3.12 Preparation of self-quenched BSA-FITC substrate

First, the accessible thiol of BSA was blocked by combining BSA (10 mg/mL) with 4-maleimidobutiric acid (1 mg/mL) in phosphate buffer (50 mM, pH 6.8) and allowing the reaction to proceed for 2 hours at room temperature. Afterwards, the reaction was purified by Amicon centrifugal filtration (MWCO 3 kDa), buffer exchanged to 10 mg/mL potassium carbonate and protein concentration set to 10 mg/mL (according to UV absorbance at 280 nm). From this solution, self-quenched BSA-FITC was prepared from a previously described protocol[3]. Briefly, FITC was added to the solution from a concentrated DMSO stock to a final concentration of 2 mg/mL (final DMSO 2 % v/v). The reaction was left at 37°C under stirring for 21 hours and purified extensively by Amicon centrifugal filtration (MWCO 3 kDa) in milliQ water until no absorbance from fluorescein could be observed in the wash waters.

##### 3.13 Activation of papain zymogens by protein activator with degradation of a protein substrate

In a 96 well plate, solution of self-quenched BSA-FITC in borate buffer (25 mM, pH 8.0) was combined with 1 μM of different zymogens ($Z_0$, $Z_{PEG}$ and $Z_{LA,}$ as determined by UV absorbance at 100 nM) and 1 μM of creatine kinase in the same buffer. Reaction volume was 100 μL and samples with DTT (1 mM) and no activator were used as positive and negative control respectively. Progress of the reaction was observed by the increase in fluorescence $\lambda_{ex}/\lambda_{em}$ 490/520 at 37° C in a plate reader for 2 h in intervals of 1 min. After 2h the plate was imaged inside an ImageQuant™ LAS 4000 camera system (GE Healthcare) using the GFP filter in fluorescene mode. These experiments were performed in 3 independent replicates (independent zymogen syntheses).

##### 3.14 Zymogen exchange experiment

In a white 96 well plate, samples containing papain inhibitor 10 µM (E-64, Sigma), 3 µM (determined by UV absorbance at 280 nm) of papain zymogens ($Z_0$, $Z_{LA}$) in Tris-HCl buffer (20 mM, pH 8.0). Then, creatine kinase (CK) was added to the wells in a concentration of 0.1 µM and incubated for 1h at room temperature. After this time, CK substrates ADP, CP and luciferin at a concentration of 1 µM and Cell-Titer Glo® reagent in a 1:10 dilution were added as described above (final volume in well 100 µL) and luminescence was monitored at 37$^o$ C in regular time intervals 1 hour (RLU integrated every 2s) in a plate reader. These experiments were performed in 3 independent replicates (independent zymogen syntheses).

#### 4. Analysis

**4.2 Statistical analysis.** Where reported, statistical significance was evaluated with a two-way ANOVA with the Sidak's multiple comparisons test performed in the software Graphpad Prism®.

**4.3 Cysteine Surface Accessibility Analysis.** The proteins tertiary structures were downloaded from the Protein Data Bank. Molecular graphics and solvent accessible surface areas (Å$^2$) calculations of cysteine residues were performed with UCSF Chimera (Resource for Biocomputing, Visualization and Informatics at the University of California, San Francisco, with support from NIH P41-GM103311). Accessible surface area values were determined with a probe radius of 1.4 Å, equivalent to the radius of a water molecule. As a reference for cysteine accessibility, calculated values were compared to reported accessible surface area of a cysteine residue in a folded protein (5 Å$^2$)[1]. In this analysis, all cysteine residues with calculated values above 5 Å$^2$ were considered accessible.
